# Population structure and antimicrobial resistance patterns of *Salmonella* Typhi isolates in Bangladesh from 2004 to 2016

**DOI:** 10.1101/664136

**Authors:** Sadia Isfat Ara Rahman, Zoe Anne Dyson, Elizabeth J. Klemm, Farhana Khanam, Kathryn E. Holt, Emran Kabir Chowdhury, Gordon Dougan, Firdausi Qadri

## Abstract

**Background:** Multi-drug resistant typhoid fever remains an enormous public health threat in low and middle-income countries. However, we still lack a detailed understanding of the epidemiology and genomics of *S.* Typhi in many regions. Here we have undertaken a detailed genomic analysis of typhoid in Bangladesh to unravel the population structure and antimicrobial resistance patterns in *S.* Typhi isolated in between 2004-2016.

**Principal findings:** Whole genome sequencing of 202 *S.* Typhi isolates obtained from three study locations in urban Dhaka revealed a diverse range of *S.* Typhi genotypes and AMR profiles. The bacterial population within Dhaka were relatively homogenous with little stratification between different healthcare facilities or age groups. We also observed evidence of transmission of Bangladeshi genotypes with neighboring South Asian countries (India, Pakistan and Nepal) suggesting these are circulating throughout the region. This analysis revealed a decline in H58 (genotype 4.3.1) isolates from 2011 onwards, coinciding with a rise in a diverse range of non-H58 genotypes and a simultaneous rise in isolates with reduced susceptibility to fluoroquinolones, potentially reflecting a change in treatment practices. We identified a novel *S.* Typhi genotype, subclade 3.3.2 (previously defined only to clade level, 3.3), which formed two localised clusters (3.3.2.Bd1 and 3.3.2.Bd2) associated with different mutations in the Quinolone Resistance Determining Region (QRDR) of gene *gyrA*.

**Significance:** Our analysis of *S*. Typhi isolates from Bangladesh isolated over a twelve year period identified a diverse range of AMR profiles and genotypes. The observed increase in non-H58 genotypes associated with reduced fluoroquinolone susceptibility may reflect a change in treatment practice in this region and highlights the importance of continued molecular surveillance to monitor the ongoing evolution of AMR in Bangladesh. We have defined new genotypes and lineages of Bangladeshi *S.* Typhi which will facilitate identification of these emerging AMR clones in future surveillance efforts.

**Author Summary:** Typhoid fever, caused by *Salmonella enterica* serovar Typhi, is an acute and often life-threatening febrile illness in developing countries. Until recently, there have been limited studies focusing on the epidemiology and disease burden of typhoid in poor resource settings including Bangladesh. This study highlights the urgent need for sustained genomics based surveillance studies to monitor the population structure and ongoing evolution of AMR. Our data revealed a diverse range of *S.* Typhi genotypes and AMR patterns among 202 isolates collected from three urban areas in Dhaka. Moreover, we defined a new genotype, subclade 3.3.2 (previously typed only to clade level, 3.3) with two Bangladesh-localiased clades 3.3.2.Bd1 and 3.3.2.Bd2 showing reduced susceptibility to fluoroquinolones. Our data shows a significant increase in non-H58 genotypes carrying QRDR mutations from 2012 onwards, replacing MDR H58 genotypes. Our data suggest that a shift in treatment practice towards third generation cephalosporins to control typhoid might be beneficial, in addition to the introduction of vaccination programs and improvements in water sanitation and hygiene (WASH) in Bangladesh.

## Introduction

*Salmonella enterica* serovar Typhi (*S*. Typhi), the causative agent of typhoid fever, is a facultative intracellular and human restricted pathogen predominantly transmitted by the fecal-oral route. Typhoid remains an enormous public health threat in many developing countries due to inadequate access to safe water, poor sanitation systems and inappropriate use of antimicrobial drugs. It is estimated that typhoid fever affects 12-27 million people globally each year whereas in Bangladesh the overall incidence is estimated at between 292-395 cases per 100,000 people per year [1–5]. Multi-drug resistant (MDR) *S*. Typhi, defined as resistance to the first-line antibiotics ampicillin, chloramphenicol and trimethoprim-sulfamethoxazole, was first observed in the 1970s [6–9]. The more recent emergence of MDR *S.* Typhi with nalidixic acid resistance and reduced susceptibility to fluoroquinolones complicates treatment options. However, third generation cephalosporins such as ceftriaxone and cefixime have proven to be effective choices for treatment as resistance to cephalosporin in *S*. Typhi is still relatively rare. The first major outbreak with ceftriaxone resistance (extensively drug resistant, defined as resistant to three first-line drugs, fluoroquinolones and third-generation cephalosporin) was observed in Pakistan from 2016 onwards [7, 10, 11].

The acquisition of antimicrobial resistance (AMR) genes by *S*. Typhi was historically associated with self-transmissible IncHI1 plasmids that harbor composite transposons [9, 12]. The global burden of MDR typhoid is driven to a significant degree by the dissemination of the highly clonal, expanding haplotype H58 (genotype 4.3.1), which is now dominant in many endemic settings throughout Africa and Asia [7, 9, 13, 14]. H58 *S.* Typhi encoding nonsynonymous mutations in the quinolone resistance determining region (QRDR) of DNA gyrase genes *gyrA* and *gyrB* and DNA topoisomerase IV genes *parC* and *parE* have shown reduced susceptibility to fluoroquinolones [7, 15]. Studies on typhoid in Nepal have reported the evolution of a fluoroquinolone resistant H58 lineage II isolates carrying three QRDR mutations (*gyrA-S83F, gyrA-D87N*, and *parC-S80I*) responsible for the failure of a gatifloxacin treatment trial [6, 15]. More recently, in Bangladesh, a new H58 lineage I triple QRDR mutant carrying three different mutations (*gyrA-S83F, gyrA-D87G*, and *parC-E84K*) has been observed [14]. This study also defined a H58 “lineage Bd” containing the sublineage “Bdq” that is characterized by a high median minimal inhibitory concentration (MIC) to ciprofloxacin (4 µg/mL) potentially involving a *qnrS* gene in addition to a *gyrA-*S83Y mutation [14].

The lack of credible surveillance data representing the actual disease burden of typhoid fever in Bangladesh presents a barrier to the implementation of control strategies. Thus, there is an urgent need for sustained genomics-based surveillance studies in poor resource settings like Bangladesh to monitor the pathogen population structure, transmission dynamics, AMR patterns and the impacts of control strategies such as vaccination programs. Here, we have used whole genome sequencing (WGS) to understand the population structure of *S*. Typhi isolated from three different urban areas of Dhaka, Bangladesh, between 2004 and 2016.

## Methods

### Ethics statement

Ethical approval was obtained from Research Review Committee (RRC) and the Ethical Review Committee (ERC) of the International Centre for Diarrhoeal Disease Research, Bangladesh (icddr,b). Informed written consent and clinical information were taken from legal guardians of child participants and adult participants.

### Study settings and blood sample collection

Dhaka is the capital city of Bangladesh and the most densely populated city with a population of over 18 million [16]. icddr,b is an international health research organization located at the Mohakhali area in Dhaka which runs two urban field sites at Kamalapur and Mirpur. Kamalapur is situated in southeast part of Dhaka, whereas Mirpur is located in northeast part of Dhaka metropolitan area. Both of these field sites are frequented by typhoid fever patients where sanitation systems are poor and access of safe drinking water is limited. This study was designed with these three urban areas inside Dhaka city: icddr,b Kamalapur field site, icddr,b Mirpur field site and icddr,b Dhaka hospital, Mohakhali. Suspected typhoid fever patients were enrolled from the three sites based on the criteria of fever at least 38°C with a minimum duration of 3 days. Blood samples (3 mL for children <5 years of age and 5 mL for others) from typhoid suspected patients were collected between 2004 to 2016 and were cultured using the automated BacT/Alert method [17, 18] for confirmation of typhoid fever.

### Bacterial isolation from blood culture

Specimens from positive blood culture bottles were sub-cultured on MacConkey agar plate and incubated at 37°C for 18-24 hours. *S.* Typhi colonie*s* were identified using standard biochemical test and slide agglutination test with *Salmonella*-specific antisera (Denka Sieken Tokyo, Japan) [17–19]. On the basis of blood culture confirmation result, we included all the available stored 202 *S.* Typhi isolated from 2004 to 2016 in this study and subjected these to whole genome sequencing analysis.

### DNA extraction and whole genome sequencing

Genomic DNA was extracted using the Wizard Genomic DNA Kit (Promega, Madison, WI, USA) according to the manufacturer’s instructions. Index-tagged paired-end Illumina sequencing libraries with an insert size of 500 bp were prepared as previously described [20] and combined into pools each containing 96 uniquely tagged libraries. WGS was performed at the Wellcome Trust Sanger Institute using the Illumina Hiseq2500 platform (Illumina, San Diego, CA, USA) to generate 125 bp paired-end reads. Sequence data quality was checked using FastQC (http://www.bioinformatics.babraham.ac.uk/projects/fastqc) to remove adapter sequences and low quality reads. Illumina sequence data was submitted to the European Nucleotide Archive and a full list of accession numbers for each isolate is summarised in **S1 Table**. Sequence data from 88 Bangladeshi *S.* Typhi from Wong *et al*. 2016 [21], and a further 528 from Tanmoy *et al.* 2018 [14], were also included for context in this study (raw sequence data are available in European Nucleotide Archive under study accessions ERP001718 and PRJEB27394, respectively).

### Read alignment and SNP analysis

*S.* Typhi Illumina reads were mapped to the CT18 (accession no. AL513382) reference chromosome sequence [22] using the RedDog mapping pipeline (v1beta.10.3; https://github.com/katholt/reddog). RedDog uses Bowtie (v2.2.9) [23] to map reads to the reference genome, and SAMtools (v1.3.1) [24, 25] to identify SNPs that have a phred quality score above 30, and to filter out those SNPs supported by less than five reads, or with 2.5x the average read depth that represent putative repeated sequences, or those that have ambiguous base calls. For each SNP that passes these criteria in any one isolate, consensus base calls for the SNP locus were extracted from all genomes mapped, with those having phred quality scores under 20 being treated as unknown alleles and represented with a gap character. These SNPs were then used to assign isolates to previously defined genotypes according to an extended *S.* Typhi genotyping framework using the GenoTyphi python script, available at https://github.com/katholt/genotyphi [6, 21]. Unique SNPs defining novel genotypes and lineages detected in the present study were manually extracted from allele tables output by RedDog using R, with SNPs responsible for non-synonymous mutations in highly conserved genes prioritised for lineage definitions.

Chromosomal SNPs with confident homozygous calls (phred score above 20) in >95% of the genomes mapped (representing a ‘soft’ core genome) were concatenated to form an alignment of alleles at 6,089 variant sites. SNPs called in prophage regions and repetitive sequences (354 kb; ~7.4% of bases in the CT18 reference chromosome, as defined previously) [21] or in recombinant regions as detected by Gubbins (v2.3.2) [26] were excluded resulting in a final alignment of 4,395 chromosomal SNPs out of a total alignment length of 4,462,203 bp for 818 Bangladeshi isolates. SNP alleles from *S.* Paratyphi A AKU1_12601 (accession no: FM2000053) were included for the purpose of outgroup rooting the phylogenetic tree.

To provide global context, 1,560 additional *S.* Typhi genomes belonging to the genotypes found in Bangladesh [6, 7, 9, 14, 15, 21, 27] were subjected to both SNP calling and genotyping as described above, resulting in an alignment of 14,852 chromosomal SNPs.

SNP distances were calculated using the dna.dist() function in the Analysis of Phylogenetics and Evolution (ape) R package [28]. Shannon diversity was calculated using the diversity() function in the R package vegan [29]. Unless otherwise stated all statistical analyses were carried out in R [30].

### Phylogenetic analysis

Maximum likelihood (ML) phylogenetic trees were inferred from the aforementioned chromosomal SNP alignments using RAxML (v8.2.8) [31]. A generalized time-reversible model and a Gamma distribution was used to model site-specific rate variation (GTR+ Γ substitution model; GTRGAMMA in RAxML) with 100 bootstrap pseudoreplicates used to assess branch support for the ML phylogeny. The resulting phylogenies were visualized and annotated using Microreact [32] and the R package ggtree [33].

### Antimicrobial resistance gene identification and plasmid replicon analysis

The mapping based allele typer SRST2 [34] was used in conjunction with the ARGannot [35] and PlasmidFinder [36] databases to detect acquired AMR genes and plasmid replicons, respectively. Where detected, isolates possessing IncHI1 plasmids were subjected to plasmid multilocus sequence typing (PMLST) [37] using SRST2 [34]. Where AMR genes were present without evidence of a known resistance plasmid replicon, raw sequence reads were assembled *de novo* using Unicycler (v0.3.0b) [38] and examined visually using the assembly graph viewer Bandage (0.8.1) [39] to inspect the composition and insertion sites of resistance-associated composite transposons. ISMapper (v2.0) [40] was run with default parameters to screen for insertion sites the transposases IS*1* (accession number J01730) and IS*26* (accession number X00011), relative to the CT18 reference sequence in order to identify the location of these in the chromosome of each Bangladeshi *S.* Typhi genome. Point mutations in the QRDR of genes *gyrA* and *parC* associated with reduced susceptibility to fluoroquinolones [15] were determined using GenoTyphi [6, 21].

## Results

### Population structure of *S.* Typhi in Bangladesh

A total collection of 202 Bangladeshi *S.* Typhi isolated between 2004 and 2016 **(S1 Table)** were subjected to Illumina whole genome sequencing, and were analysed together with 616 additional Bangladeshi *S.* Typhi whole genome sequences from previous studies (isolated between 1998 and 2016, see **S2 Table**) to provide a robust genomic context. The *S*. Typhi genomes were subjected to phylogenetic and GenoTyphi analysis, revealing that the pathogen population structure in Bangladesh is diverse with 17 distinct genotypes as summarised in **S3 Table** and Fig. 1. Eight *S*. Typhi subclades (genotypes 2.0.1, 2.1.7, 2.3.3, 3.0.1, 3.1.2, 3.2.2, 3.3.0, 4.3.1) and two undifferentiated isolates of major lineage 2 (genotype 2.0.0) were observed among our samples collected in Bangladesh between 2004 and 2016 (Table 1). The majority of our isolates (n=83, 41.1%) belonged to genotype 4.3.1 (haplotype H58), which has rapidly expanded in South Asia from the early 1990s. Previously defined major sublineages of H58 (lineage I and lineage II) [9] were present among the Bangladeshi isolates, with H58 lineage I (genotype 4.3.1.1) isolates appearing dominant (n=63, 31.2%). Only a single H58 lineage II (genotype 4.3.1.2) isolate (n=1, 0.50%) was present; this is somewhat surprising as this lineage is common in neighboring countries Nepal and India [6, 15]. In addition to H58 lineage I and II isolates, 19 genetically undifferentiated H58 isolates (genotype 4.3.1 according to the framework of Wong *et al* 2016) were also observed. Further analysis of these revealed close clustering with 119 “H58 lineage Bd” isolates reported by Tanmoy *et al*. 2018 [14], forming a monophyletic sister clade to H58 lineages I and II that we herein define as 4.3.1.3 (see Fig. 1). The 19 4.3.1.3 isolates from our collection possess the characteristic SNPs STY0054-C711A (position 561056 in CT18) and STY2973-A1421C (position 2849843 in CT18) that Tanmoy *et al* used to define this lineage [14]. We have therefore added the SNP STY0054-C711A to the GenoTyphi script (available at: http://github.com/katholt/genotyphi), to facilitate the detection of this lineage (genotype 4.3.1.3) in future WGS surveillance studies.

**Table 1:** Genotypes present among 202 novel *S.* Typhi isolates from Dhaka, Bangladesh. *Novel genotypes described in this manuscript.

| Genotype | No. of isolates (%) |
| --- | --- |
| 2.0.0 | 2 (0.99%) |
| 2.0.1 | 10 (4.95%) |
| 2.1.7 | 2 (0.99%) |
| 2.3.3 | 30 (14.85%) |
| 3.0.1 | 1 (0.50%) |
| 3.1.2 | 2 (0.99%) |
| 3.2.2 | 44 (21.80%) |
| 3.3.2* | 28 (13.86%) |
| 4.3.1.3* | 19 (9.41%) |
| 4.3.1.1 | 63 (31.19%) |
| 4.3.1.2 | 1 (0.50%) |

**Fig. 1.**
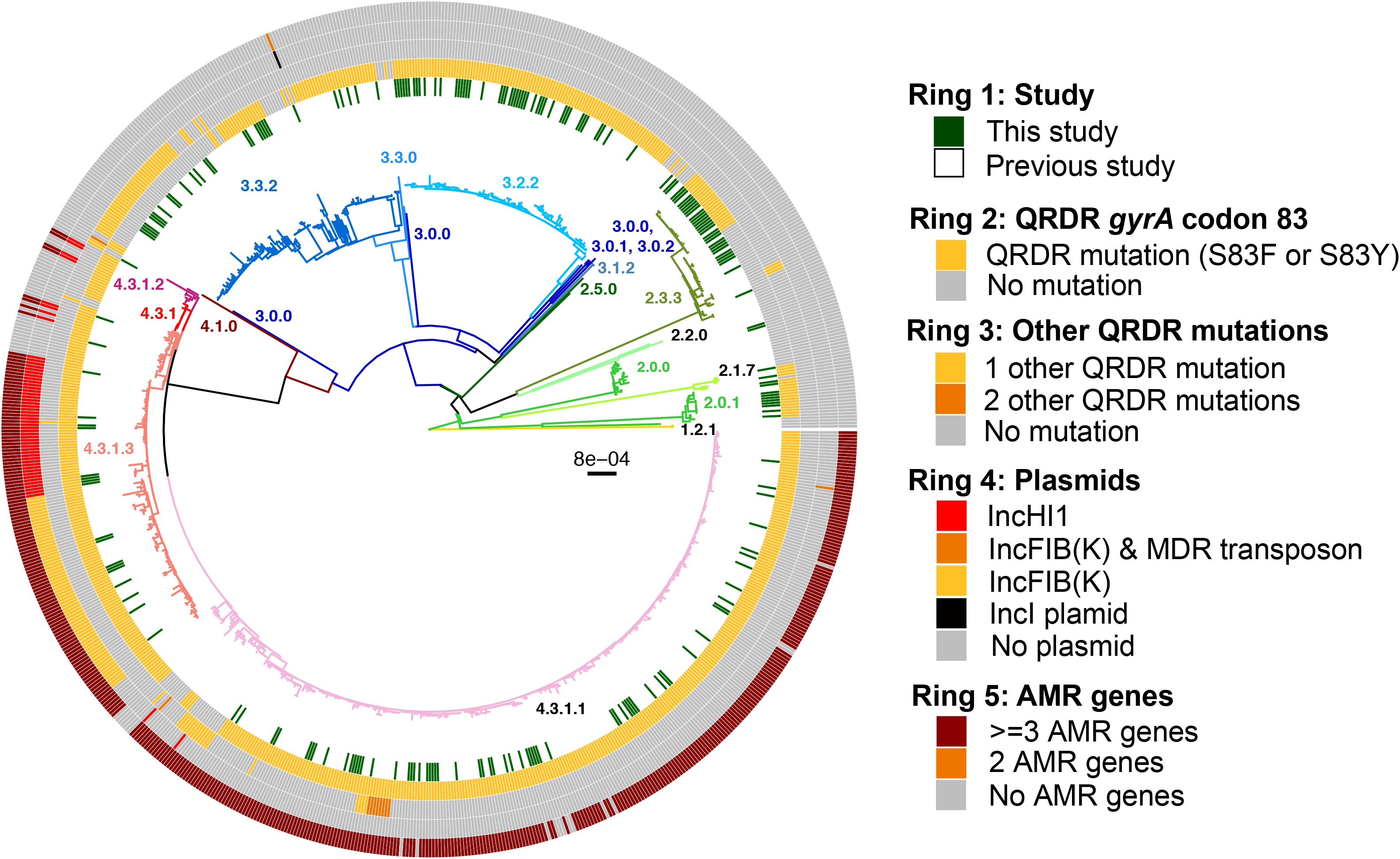
Bangladesh *S.* Typhi population structure with antimicrobial resistance. Maximum likelihood outgroup-rooted tree of 816 Bangladeshi *S.* Typhi isolates. Branch colours indicate the genotype (as labelled). Inner ring indicates the position of the 202 Bangladesh *S.* Typhi from this study, the second and third rings indicate the presence of QRDR mutations, fourth ring indicates plasmids and the outer fifth ring indicates AMR genes coloured as per the inset legend.

Genotypes 2.3.3 (n=30, 14.85%), 3.2.2 (n=44, 21.80%), 3.3.0 (n=28, 13.86%) were also relatively common in Bangladesh. Of the 818 Bangladeshi *S*. Typhi analysed, 119 (14.5%) formed a monophyletic sublineage within clade 3.3 that did not belong to any of the previously defined subclades (i.e. assigned genotype 3.3.0 in the existing scheme) and were closely related to other clade 3.3 isolates found in Nepal (median distance of ~70 SNPs) [6, 15] (see **S2 Fig** and interactive phylogeny available at http://microreact.org/project/5GzJ7Umoz). We herein assign these Bangladesh and Nepal *S*. Typhi to a novel genotype, subclade 3.3.2 (labeled in Fig. 1), which can be identified by the presence of a marker SNP STY3641-A224G (position 3498544 in CT18) that confers an amino acid mutation (Q75G) within the ST3641 encoded protein. This genotype has also been added to the GenoTyphi script to facilitate detection of subclade 3.2.2 from WGS data in future studies.

### Intra-country transmission dynamics within Bangladesh

Geographical location data were available for the 202 novel *S*. Typhi, which were collected from three different sites inside Dhaka (icddr,b Kamalapur field site, icddr,b Mirpur field site and icddr,b Dhaka hospital Mohakhali). Detailed genotypic distribution of 202 isolates from these three study sites are shown in Fig. 2, and an interactive version of the phylogeny and map are available online at https://microreact.org/project/sP2Uwk_DI). Our data showed that genotypes 2.3.3, 3.3.2, 3.2.2, 4.3.1.1 and 4.3.1.3 were present in all three study sites (Fig. 2A) and were intermingled in the phylogeny (Fig. 2B) suggesting circulation across the city. Two genotypes, 2.0.1 and 2.1.7, were restricted to two of the three study locations (Kamalapur and icddr,b hospital, and Mirpur and icddr,b hospital, respectively); genotype 2.0.0, 3.1.2 and genotype 3.0.1, 4.3.1.2 were found only in Kamalapur and icddr,b hospital, respectively.

**Fig. 2.**
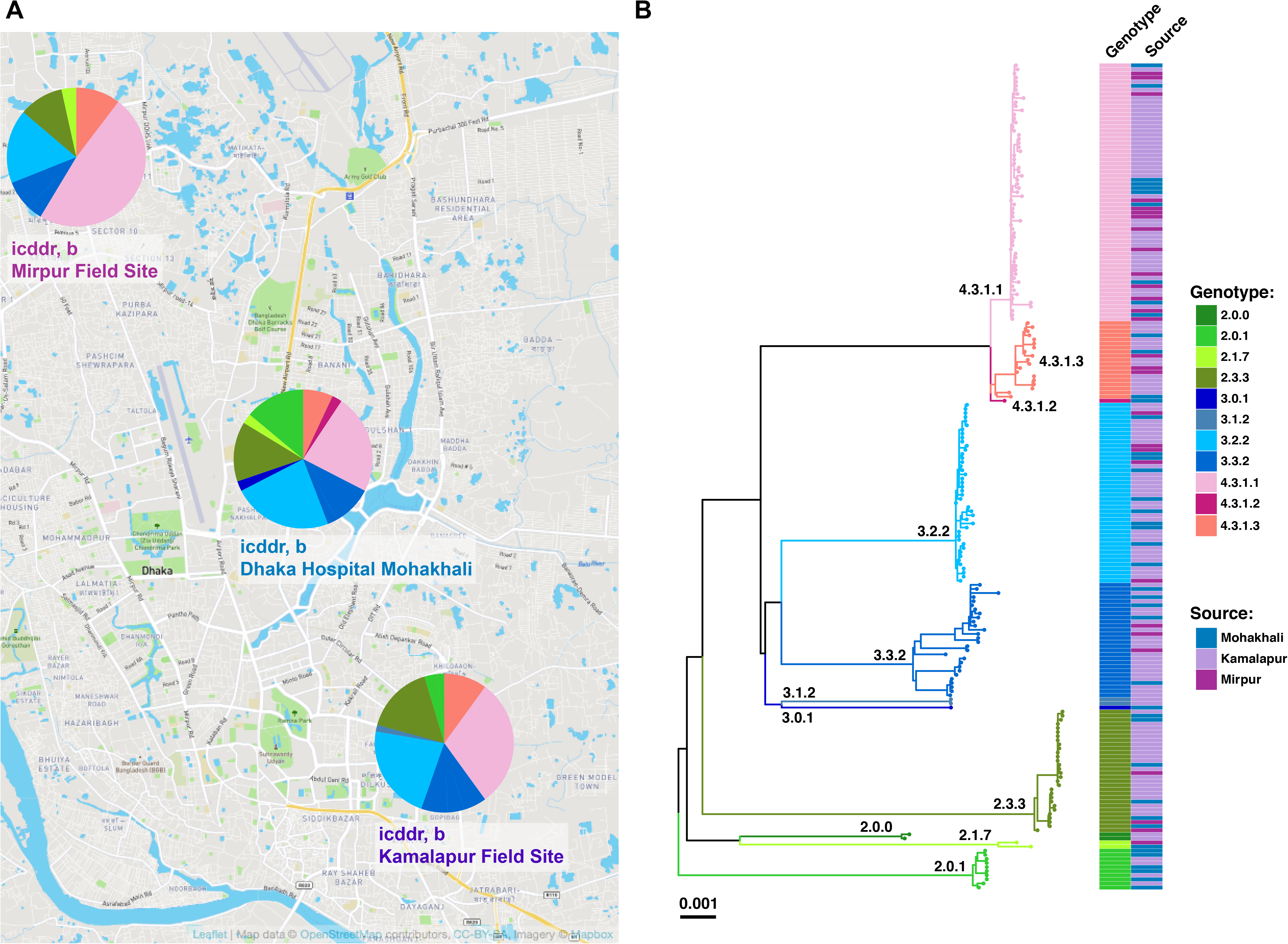
Spatial analysis of 202 *S.* Typhi from this study. **(A)** Map of Dhaka indicating the prevalence of *S.* Typhi genotypes present at each of the three study sites of icddr,b. **(B)** Maximum likelihood tree of 202 Bangladeshi *S.* Typhi isolates from the three study sites. Branch colors indicate the genotype (as labelled) and the colored heatmap (on the right) shows, for each isolate, its genotype, and source of the isolates coloured as per the inset legend.

Information on patient ages were available for 185 (91.6%) of the 202 *S.* Typhi, facilitating stratification by age groups (young children under 5 years of age, older children from 5 to 15 years of age, and adults above 15 years of age). This stratification did not reveal any significant differences (p=0.344 using Chi-squared test) as all age groups appeared to be infected with a diverse range of *S.* Typhi genotypes (Fig. 3).

**Fig. 3.**
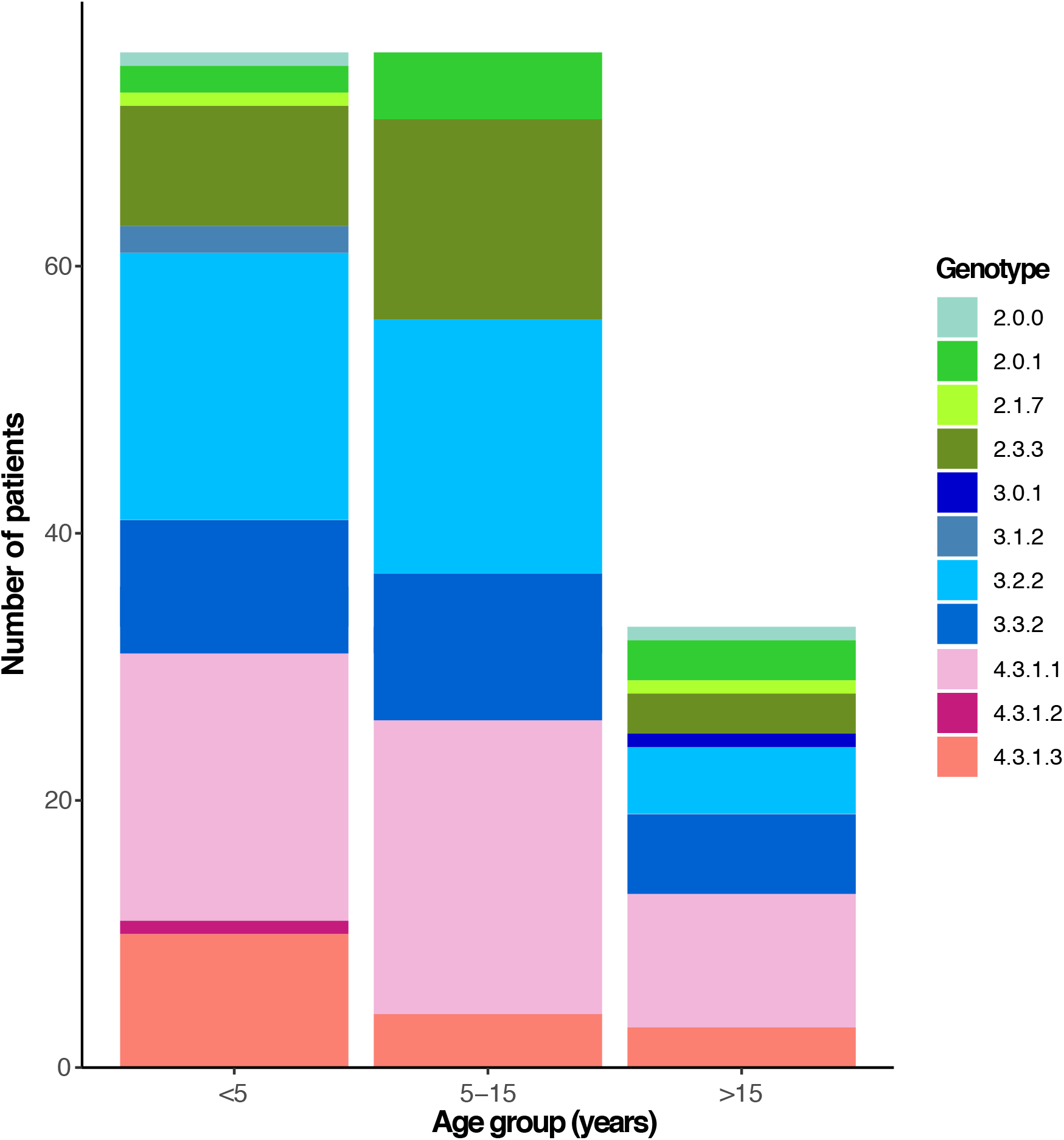
Bangladesh genotypes observed in pediatric and adult patients. Individual *S.* Typhi genotypes are colored as described in the inset legend.

### Inter-country transmission patterns and population structure of Bangladeshi *S*. Typhi

To provide a global context for the 818 Bangladeshi *S*. Typhi and to better understand inter-country transmission patterns, we constructed a global phylogeny including an additional 1,560 *S*. Typhi from over 30 countries [6, 7, 14, 15, 21, 27] from the global WGS collection, belonging to the 17 genotypes that were detected in Bangladesh (Fig. 4, interactive phylogeny available at https://microreact.org/project/5GzJ7Umoz). A single South African *S*. Typhi isolated in 2010 was intermingled among the genotype 4.3.1.3 isolates, ~2 SNPs away from its closest Bangladesh relative, suggesting that the H58 lineage Bd has been transferred from South Asia to Africa on at least one occasion. Predominantly, the Bangladeshi H58 and non-H58 *S*. Typhi were related to isolates from India, Pakistan, and Nepal (see Fig. 4 and interactive phylogeny at https://microreact.org/project/5GzJ7Umoz), suggesting regional circulation of these genotypes throughout South Asia. Notably, we found that Bangladesh isolates formed several unique monophyletic lineages within this tree, consistent with local establishment and ongoing clonal transmission within Bangladesh; e.g. 2.3.3, 3.2.2, 3.3.2 (Fig. 4A) and 4.3.1.1, 4.3.1.3 (Fig. 4B).

**Fig. 4.**
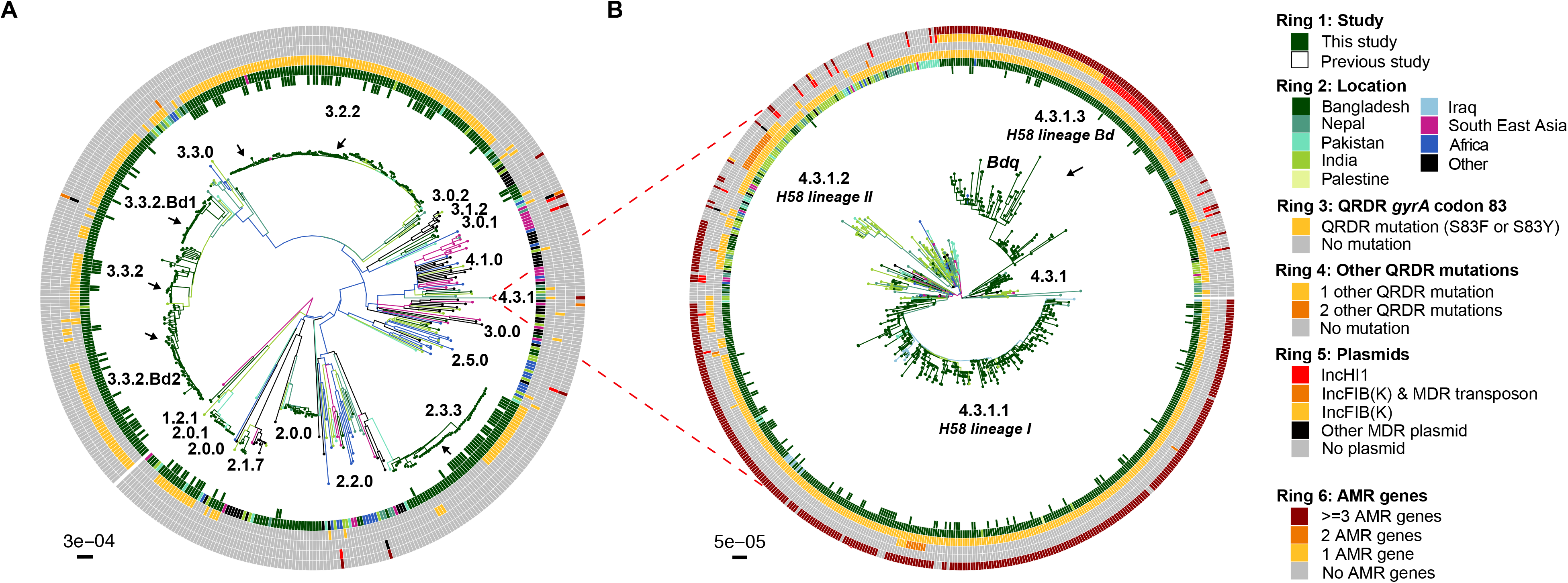
Global population structure of Bangladesh *S.* Typhi. (A) Global population structure of Non-H58 (4.3.1) Bangladesh genotypes. (B) Global population structure of H58. Branch colours indicate country/region of origin, as do the inner two rings. The third ring indicates mutations in codon 83 of gene *gyrA*, fourth ring indicate the number of additional QRDR mutations. The fourth ring indicates the presence of any plasmids and the sixth ring indicates the presence of AMR genes. All branches and rings are coloured as per the inset legend. Arrows indicate localised lineages of Bangladeshi isolates.

### Antimicrobial resistance and plasmid replicons in *S.* Typhi in Bangladesh

To better understand the AMR burden among *S.* Typhi in Bangladesh, we subjected our 202 isolates (Table 2) and the additional 616 *S*. Typhi from previous studies [9, 14, 21] (**S4 Table**) to screening for both genes and mutations associated with AMR. Only 17 of our S. Typhi (9.42%) lacked any known molecular determinants of AMR and were thus predicted to be fully susceptible to antibiotics (Fig. 1 and Table 2).

**Table 2:** Genetic determinants of antimicrobial resistance in 202 S. Typhi isolates from Dhaka, Bangladesh

| Resistance patterns | H58 isolates<br>(n=83) | Non H58<br>isolates<br>(n=119) | Total<br>isolates<br>(n=202) |
| --- | --- | --- | --- |
| **Acquired AMR genes / QRDR mutations** | **74 (89.16%)** | **0 (0.00%)** | **74 (36.63%)** |
| *bla*<sub>TEM-1</sub>, *catA1*, *dfrA7*, *sul1*, *sul2*, *strAB* / *gyrA*-S83F | 47 (56.63%) | 0 (0.00%) | 47 (23.27%) |
| *bla*<sub>TEM-1</sub>, *catA1*, *dfrA7*, *sul1*, *sul2*, *strAB*, *tetB* / *gyrA*-S83Y | 8 (9.64%) | 0 (0.00%) | 8 (3.96%) |
| *bla*<sub>TEM-1</sub>, *catA1*, *dfrA7*, *sul1*, *sul2*, *strAB* / *gyrA*-S83F, *parC*-E84K | 1 (1.20%) | 0 (0.00%) | 1 (0.50%) |
| *bla*<sub>TEM-1</sub>, *catA1*, *dfrA7*, *sul1* / *gyrA*-S83F | 1 (1.20%) | 0 (0.00%) | 1 (0.50%) |
| *catA1*, *dfrA7*, *sul1* / *gyrA*-S83F | 8 (9.64%) | 0 (0.00%) | 8 (3.96%) |
| *bla*<sub>TEM-1</sub>, *sul2*, *qnrS*, *tetA* / *gyrA*-S83Y | 9 (10.84%) | 0 (0.00%) | 9 (4.46%) |
| **QRDR mutations only** | **83 (100%)** | **102 (85.71%)** | **185 (91.6%)** |
| *gyrA*-S83F | 62 (74.70%) | 85 (83.33%) | 147 (72.77%) |
| *gyrA*-S83Y | 19 (22.90%) | 3 (2.94%) | 22 (10.89%) |
| *gyrA*-D87G | 1 (1.20%) | 0 (0.00%) | 1 (0.50%) |
| *gyrA*-D87N | 0 (0.00%) | 12 (11.76%) | 12 (5.94%) |
| *gyrA*-D87Y | 0 (0.00%) | 2 (1.96%) | 2 (0.99%) |
| *gyrA*-S83F, *parC*-E84K | 1 (1.20%) | 0 (0.00%) | 1 (0.50%) |
| **Susceptible to all antibiotics** | **0 (0.00%)** | **17 (16.67%)** | **17 (8.42%)** |

Many H58 isolates (n=57, 68.67%) were predicted to be MDR, carrying genes associated with resistance to the first-line drugs chloramphenicol, trimethoprim-sulfomethoxazole and ampicillin. The majority of these were H58 lineage I isolates (genotype 4.3.1.1, n=48, 64.86%) carrying genes *catA1, dfrA7, sul1, sul2, bla*_TEM-1_ and *strAB* (**S1 Fig**) in a composite transposon (Fig. 5) conferring resistance to all three first-line drugs and also streptomycin. These isolates lacked the IncHI1 plasmid, suggesting that the AMR genes have been integrated into the chromosome. A small proportion of genotype 4.3.1.1 *S*. Typhi (n=8, 10.8%) carried an alternative form of the typical transposon encoding just three AMR genes (*catA1, dfrA7, sul1*; see Fig. 5). Examination of the assembly graphs and nucleotide sequence comparisons [41] of the genomes carrying the 3-gene vs 7-gene locus revealed that integration of both transposons were mediated by IS*1* transposition associated with an 8 bp target site duplication (GGTTTAGA; see Fig 5). However as IS*1* was present at multiple locations in the chromosome sequences of these isolates, we were unable to resolve the precise location of the MDR integration site, and insertions at either of two previously reported chromosomal integration sites (near *cyaA* or within *yidA* [9]) were equally possible in the Bangladeshi isolates.

**Fig. 5.**
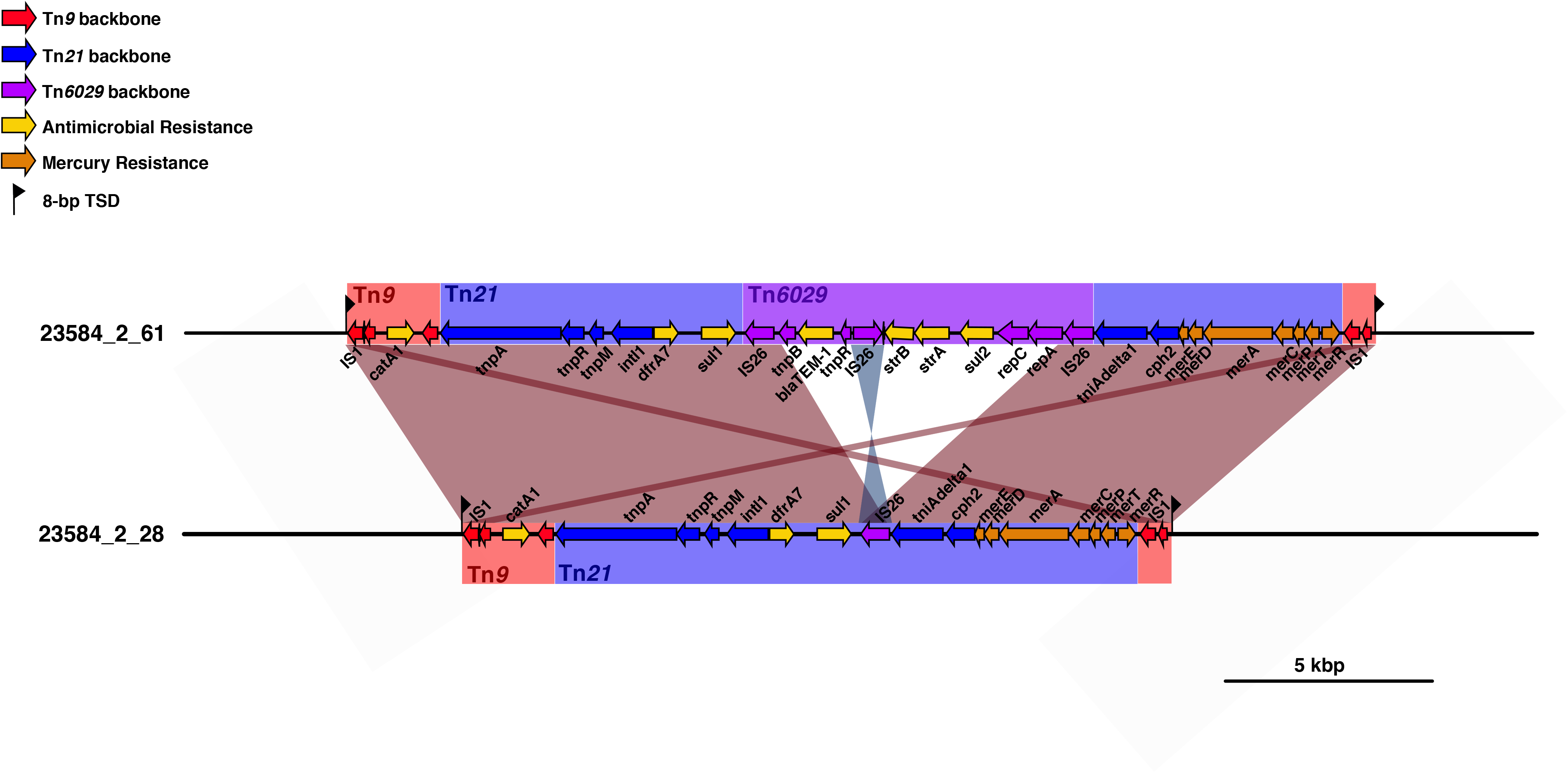
Insertion sites of transposons observed in *S.* Typhi from Bangladesh. Genes and transposons are indicated according to the inset legend. TSD indicates target site duplication, and *Tn* indicates transposons.

Of the 19 H58 lineage Bd isolates detected in our collection, two different plasmid mediated AMR patterns emerged (**Fig. S2**). The first pattern (n=8) was characterized by the presence of the IncHI1 plasmid (plasmid sequence type, PST6) [9] carrying eight AMR genes (*bla*_*TEM-1*_, *catA1, dfrA7, sul1, sul2, strAB*, and *tetB*) conferring resistance to the first line drugs plus streptomycin and tetracyclines. The second pattern (n=9) was characterized by the presence of an IncFIB(K) plasmid carrying the AMR genes (*bla*_*TEM-1*_, *sul2, tetA*) conferring resistance to ampicillin, sulfonamides, tetracyclines, and also *qnrS* together with the *gyrA*-S83Y mutation confers resistance to fluoroquinolones. These isolates were intermingled with IncFIB(K)-carrying isolates described by Tanmoy *et al* 2018 (which they termed “sublineage Bdq”). This IncFIB(K)-carrying cluster appears to have emerged from the main 4.3.1.3 group that typically carries the IncHI1 plasmid (separated by a median of ~11 SNPs), but the IncHI1 MDR plasmid has been replaced in this group by the IncFIB(K) fluoroquinolone resistance plasmid (see **S2 Figure**). The IncFIB(K)-containing Bdq cluster appears to have undergone a clonal expansion in Bangladesh, but has not replaced the IncHI1 form of 4.3.1.3 (see below).

Overall, non-synonymous mutations in the QRDR of genes *gyrA* and *parC* associated with reduced susceptibility to fluoroquinolones (FQ) were common among our Bangladeshi *S.* Typhi isolates (n=185, 91.6%) harboring at least one QRDR mutation (Table 2). Unlike the acquisition of MDR genes, the QRDR mutations (mainly in gene *gyrA* at codon 83) were common in both non-H58 (n=102, 85.71%) as well as H58 isolates (n=83, 100%). Examination of genotype 3.3.2 revealed two distinct monophyletic lineages of Bangladeshi *S*. Typhi of this genotype each with a different QRDR mutation. Here we defined these two Bangladeshi lineages as 3.3.2.Bd1 and 3.3.2.Bd2, carrying the *gyrA-*S83F and *gyrA*-D87N mutations, respectively (Fig. 4A). Markers for these two lineages (SNPs STY2588-G378A, position 2424394 in CT18; and STY2441-G439A, position 2272144 in CT18; respectively) have been added to the GenoTyphi script to facilitate their detection. No QRDR triple mutants were observed among our collection; however, a single double mutant *S*. Typhi of genotype 4.3.1.1 was identified carrying both *gyrA*-S83F and *parC-*E84K mutations. Hence, the only isolates we predict to be ciprofloxacin resistant are the IncFIB(K) group carrying *qnrS* and *gyrA*-S83Y.

### Temporal trends in genotypic distribution and AMR over time

We examined the genotypic distribution and AMR patterns over time during our study period of 2004 to 2016; note our sampling includes all available blood culture positive *S*. Typhi from the three sites in urban Dhaka. Prior to 2011, H58 lineages (4.3.1.1 and 4.3.1.3) dominated the population across the three sites sampled (median 66.67% per year; see Fig. 6A). During this time most isolates were MDR (median 55.56% per year, see Fig. 6B). After this period, there appears to be a relative decrease in the frequency of H58 genotypes (4.3.1.1, 4.3.1.2, 4.3.1.3; median 28.03% per year in 2011-2016) and an increase in non-H58 genotypes 2.3.3, 3.2.2, 3.3.2 (from median 22.2% per year in 2004-2010 to 65.2% per year in 2011-2016; overall diversity increasing from 1.34 Shannon diversity in 2004-2010 to 1.83 in 2011-2016). The decline in non-H58 *S*. Typhi coincided with a decline in MDR (median 11.88% per year in 2011-2016), with the most common profile being non-MDR with a single QRDR mutation (*gyrA-*S83F being the most prevalent). Notably, the IncFIB(K)/*qnrS* lineage Bdq remained at low frequency throughout the study period (detected from 2007 to 2014, median 4.78% per year).

**Fig 6.**
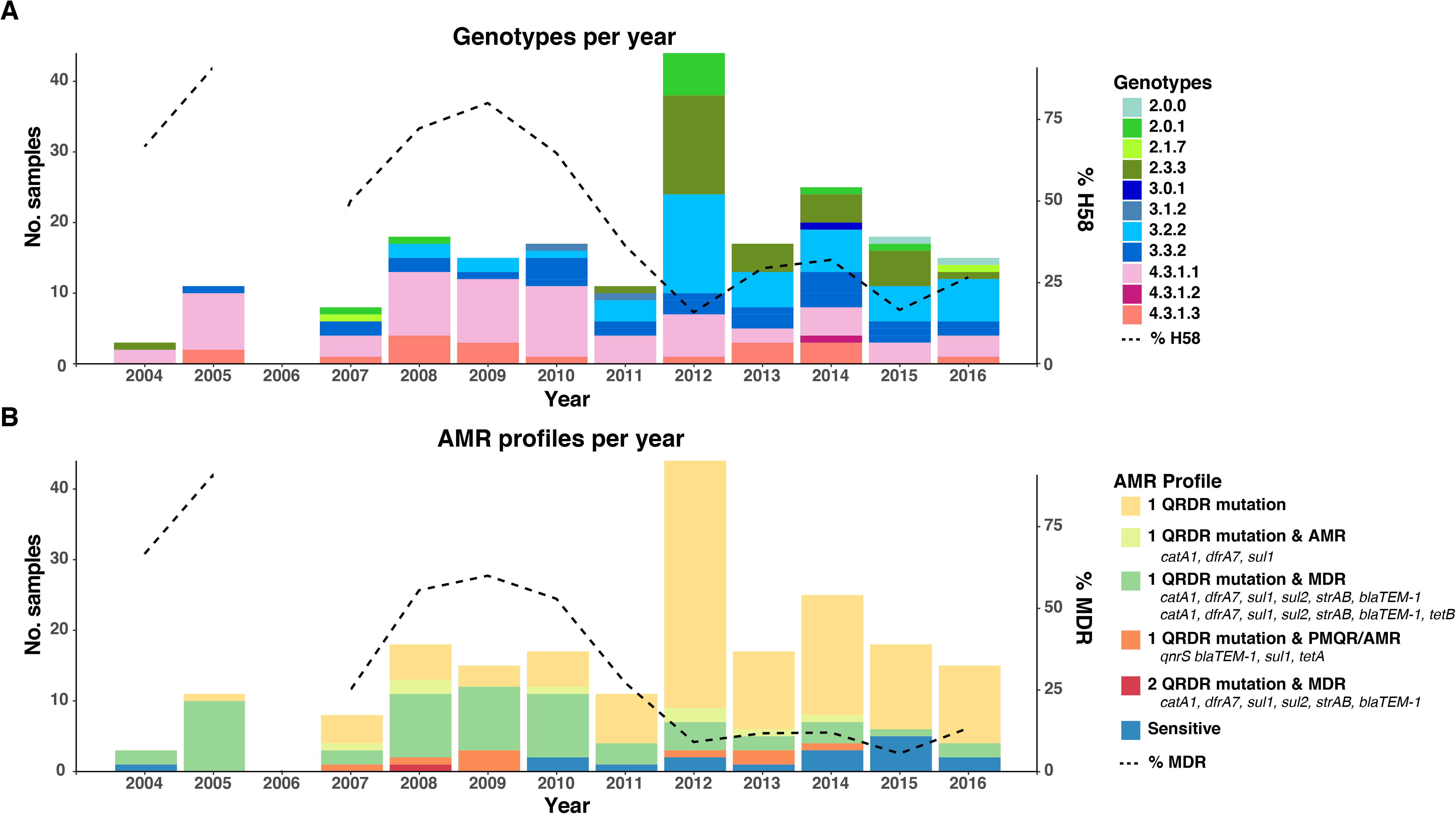
Timeline of Bangladesh *S.* Typhi genotypes and AMR profiles from 2004-2016. (A) Genotypes observed per annum. Overlaid line indicates the percentage of H58 isolates per year. (B) AMR profiles observed per annum. Overlaid line indicates the percentage of MDR isolates per year. Genotypes and AMR profiles coloured as per inset legend.

## Discussion

We show here that the population structure of *S*. Typhi in Bangladesh is diverse, harboring 17 distinct genotypes, with 9 genotypes circulating within urban Dhaka between 2004-2016 (Fig 1). There was little evidence of any local geographic restriction of *S*. Typhi genotypes within the city of Dhaka (Fig 2) and no obvious stratification of *S.* Typhi genotypes by patient age (Fig 3), consistent with what has been observed in other South Asian settings [6, 15, 42].

*S*. Typhi circulating in Bangladesh are closely related to isolates from neighboring India, Pakistan and Nepal suggesting circulation throughout South Asia. However, the formation of multiple localised lineages indicates establishment and ongoing local transmission of multiple genotypes in parallel within Bangladesh; and warranted definition of novel Bangladesh-specific subclades for future tracking via WGS surveillance. Firstly, the most common genotype in our collection was H58 lineage I (31.2%) and H58 lineage II was rare (0.5%), in contrast to neighboring India and Nepal where lineage II are highly prevalent and lineage I is rarely reported [6, 9, 15]. Secondly, this distinction between Bangladeshi and Indian/Nepali pathogen populations was also evident for the newly defined H58 lineage Bd (4.3.1.3) [14], which so far has been almost exclusively found in Bangladesh (the exceptions being singleton isolates reported in Nepal and South Africa). Thirdly, we identified a novel subclade of *S*. Typhi, genotype 3.3.2, which included a Bangladesh-specific monophyletic group with relatives in Nepal, that we further divided into 3.3.2.Bd1 and 3.3.2.Bd2 based on distinct QRDR mutations conferring reduced susceptibility to fluoroquinolones (Fig 4A). These novel Bangladesh-associated lineages (4.3.1.3, 3.3.2.Bd1, 3.3.2.Bd2) have been added to the GenoTyphi genotyping scheme, which will facilitate their detection and tracking in future surveillance efforts, and over time will reveal whether they remain localized to Bangladesh or being to disseminate through Asia and Africa as has been observed for H58 lineages I and II [6, 13, 27, 43].

The sustained, very high frequency of *S*. Typhi carrying mutations associated with reduced susceptibility to fluoroquinolones that we detected in this study (>66% per year; **see** Fig 6B) is likely the result of an increase in over-the-counter sale (without prescription) of this antibiotic class over the last decade [44]. This, along with a decrease in MDR is similar to reports from both India and Nepal [6, 9]. However, while the prevalence of reduced susceptibility was very high, we found limited evidence of evolution towards full resistance, with the *qnrS*-positive clade (associated with ciprofloxacin MIC of 4 µg/mL, [14]) remaining at low frequency throughout the study period. An H58 lineage I (4.3.1.1) QRDR triple mutant has been previously reported in Bangladesh [14], however, this was not observed among our collection; we only detected a single QRDR double mutant in H58 lineage I in 2008 (see Table 2 and Fig. 6).

Notably, the reduced prevalence of MDR coincided with a reduction in H58 (4.3.1) isolates across our 3 study sites and a significant diversification in the pathogen population, particularly driven by increased prevalence of QRDR single-mutant *S.* Typhi genotypes 2.3.3, 3.2.2 and 3.3.2. This unexpected change in population structure that cannot be explained by selection for AMR suggests unknown selective pressures may be influencing the pathogen population in Bangladesh, and highlights the need for ongoing genomic surveillance. Further, while MDR was less frequent after 2010, the presence of multiple forms of the MDR chromosomal insertion is highly concerning, as such insertions may facilitate more stable transmission of the MDR phenotype [6, 9]. Similarly, the persistent presence of plasmid-mediated quinolone resistance (PMQR) via an IncFIB(K) plasmid carrying a *qnrS1* gene in Bangladesh is concerning, despite the relatively low frequency (4.5%) at which it is observed currently.

### Conclusion

This study demonstrates the importance of molecular based surveillance studies in endemic regions, especially in Bangladesh, where the disease burden is high and many different AMR phenotypes were observed. The change in both population structure and AMR patterns over twelve years (2004 to 2012) shows increased prevalence of populations with reduced susceptibility fluoroquinolones, emphasizing the ongoing evolution of AMR in this setting as well as the urgent need for WGS based surveillance in Bangladesh to inform both treatment guidelines and control strategies.

## Acknowledgements

This work was supported by the Wellcome Sanger Institute, Cambridge, UK and International Centre for Diarrhoeal Disease Research, Bangladesh (icddr,b). This study was supported by the grants from the Wellcome Trust, the National Institutes of Health, including the National Institute of Allergy and Infectious Diseases (AI100023 [FQ]); AI058935 [FQ]) as well as a Fogarty International Center Training Grant in Vaccine Development and Public Health (TW005572 [FK and FQ]), Bill and Melinda Gates Foundation (Grant no. OPP50419) and also SIDA fund (54100020, 51060029). ZAD was supported by a grant funded by the Wellcome Trust (106158/Z/14/Z). KEH is supported by a Senior Medical Resarch Fellowship from the Viertel Foundation of Australia. We acknowledge the support of dedicated field and laboratory workers at the icddr,b involved in this study. We would like to thank Doli Goswami, Md. Lokman Hossain and Abdullah Brooks for their contribution in Kamalapur field study. icddr,b is grateful to the Governments of Bangladesh, Canada, Sweden and UK for providing core support. We would also like to thank the members of Pathogen Informatics Team and core sequencing teams at the Wellcome Sanger Institute.

## Author contributions

GD, FQ, ZAD, SIAR contributed to the design of the study and GD, FQ supervised the study. SIAR, ZAD, EJK and KEH performed the genomic data analysis. SIAR, ZAD, KEH, FQ and GD wrote the manuscript. All authors contributed to the interpretation of results, editing of the manuscript and approved the final version.

## Supporting Information

**S1 Fig. Bangladesh *S.* Typhi population structure showing acquired AMR genes, and plasmids.** Branches are coloured by genotype as labelled, the heatmap the molecular determinants of antimicrobial resistance and the presence of plasmids coloured as per the inset legend.

**S2 Fig. Population structure and country of origin of genotype 3.3.2 isolates.** Maximum likelihood phylogeny of genotype 3.3.2 isolates. Coloured bar indicates country of origin as per the inset legend.

**S1 Table. Data of 202 S. Typhi isolates from 3 different study sites in Dhaka, Bangladesh between 2004-2016**

**S2 Table. Data of 818 S. Typhi isolates from Bangladesh including previous studies between 1998-2016**

**S3 Table. Genotyping results for the 818 S. Typhi isolates from Bangladesh.**

**S4 Table. Genetic determinants of antimicrobial resistance in S. Typhi isolates from Bangladesh**

