## Supplementary figures and images for "Population structure and antimicrobial resistance patterns of *Salmonella* Typhi isolates in Bangladesh from 2004 to 2016"

### S1 Fig

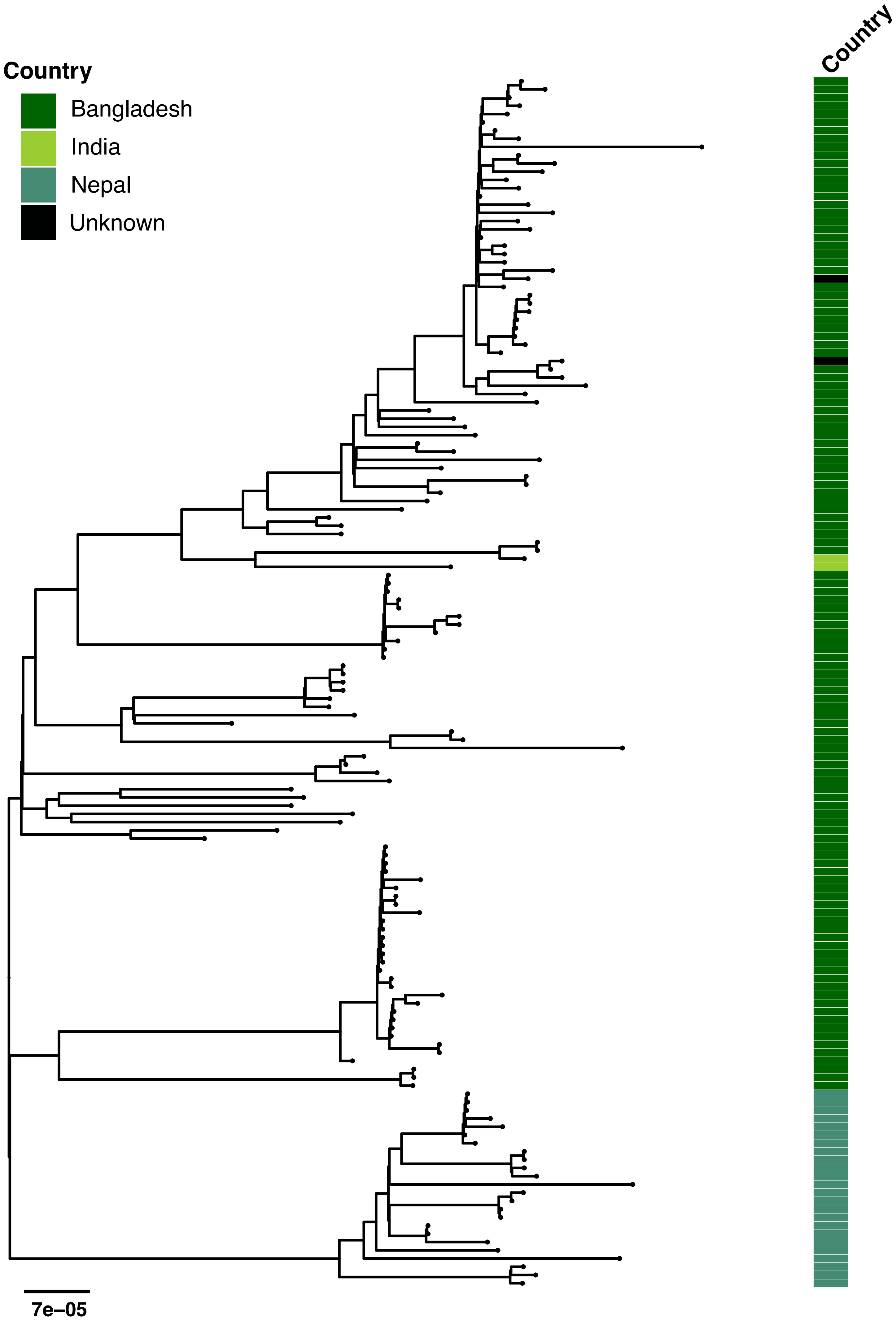

### S2 Fig

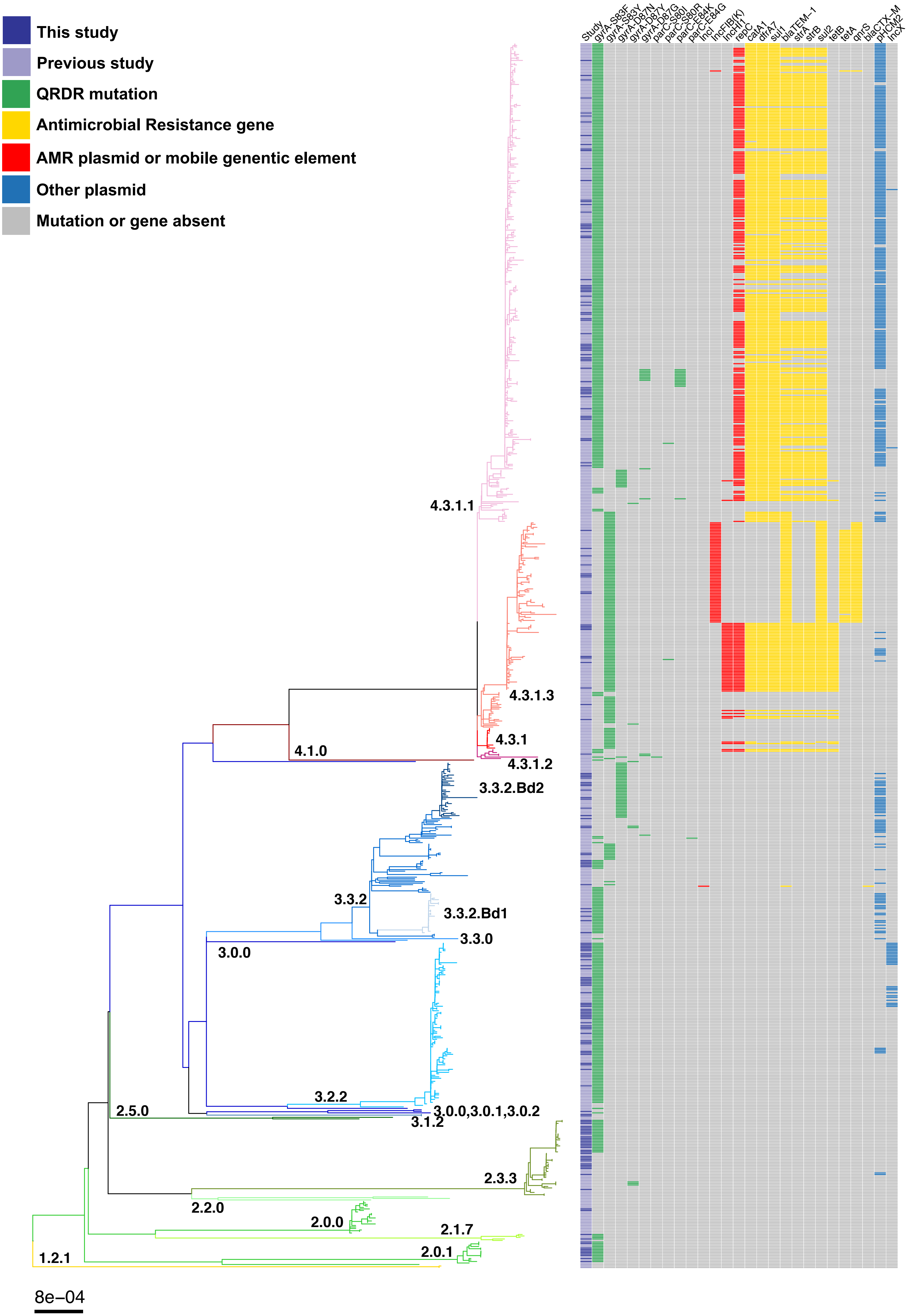
