## Supplementary material for "Population structure and antimicrobial resistance patterns of *Salmonella* Typhi isolates in Bangladesh from 2004 to 2016": S3_S4 Table

**Table S3.** Genotypes of 818 *S.* Typhi isolates from Bangladesh.

| **Genotype** | **No. of isolate (%)** |
| --- | --- |
| 1.2.1 | 2 (0.25%) |
| 2.0.0 | 23 (2.81%) |
| 2.0.1 | 15 (1.83%) |
| 2.1.7 | 5 (0.61%) |
| 2.2.0 | 3 (0.37%) |
| 2.3.3 | 51 (6.24%) |
| 2.5.0 | 2 (0.25%) |
| 3.0.0 | 2 (0.25%) |
| 3.0.1 | 3 (0.37%) |
| 3.0.2 | 1 (0.12%) |
| 3.1.2 | 2 (0.25%) |
| 3.2.2 | 110 (13.5%) |
| 3.3.0 | 2 (0.24%) |
| 3.3.2 | 117 (14.3%) |
| 4.1.0 | 1 (0.12%) |
| 4.3.1 | 14 (1.71%) |
| 4.3.1.Bd | 138 (16.9%) |
| 4.3.1.1 | 320 (39.1%) |
| 4.3.1.2 | 7 (0.86%) |

**Table S4.** Genetic determinants of antimicrobial resistance in 818 Bangladeshi *S.* Typhi isolates

| **Resistance patterns** | **H58 isolates (n=479)** | **Non H58 isolates (n=339)** | **Total isolates (n=818)** |
| --- | --- | --- | --- |
| **Acquired AMR genes** | **422 (88.1%)** | **1 (0.29%)** | **422 (51.6%)** |
| *bla_TEM-1_* *, catA1, dfrA7, sul1, sul2, strAB* / *gyrA-*S83F | 211 (44.1%) | 0 (0.00%) | 211 (25.79%) |
| *bla_TEM-1_* *, catA1, dfrA7, sul1, sul2, strAB* / *gyrA-*S83Y | 1 (0.21%) | 0 (0.00%) | 1 (0.12%) |
| *bla_TEM-1_* *, catA1, dfrA7, sul1, sul2, strA*B / *gyrA-*D87N | 9 (1.9%) | 0 (0.00%) | 9 (1.10%) |
| *bla_TEM-1_* *, catA1, dfrA7, sul1, sul2, strAB /* *gyrA-*D87G | 1 (0.21%) | 0 (0.00%) | 1 (0.12%) |
| *bla_TEM-1_* *, catA1, dfrA7, sul1, sul2, strAB /* *gyrA-*S83F, *parC-*E84K | 4 (0.84%) | 0 (0.00%) | 4 (0.49%) |
| *bla_TEM-1_* *, catA1, dfrA7, sul1, sul2, strAB /* *gyrA-*S83F, *parC-*S80R | 1 (0.21%) | 0 (0.00%) | 1 (0.12%) |
| *bla_TEM-1_* *, catA1, dfrA7, sul1, sul2, strA, strB /* *gyrA* D87G, *parC* E84K | 1 (0.21%) | 0 (0.00%) | 1 (0.12%) |
| *bla_TEM-1_* *, catA1, dfrA7, sul1, sul2, strAB /* *gyrA-*S83F, *gyrA-*D87G, *parC-*E84K | 7 (1.5%) | 0 (0.00%) | 7 (0.86%) |
| *bla_TEM-1_, catA1, dfrA7, sul1, sul2, strAB* | 2 (0.42%) | 0 (0.00%) | 2 (0.25%) |
| *bla_TEM-1_* *, catA1, dfrA7, sul1, sul2, strAB, tetB* / *gyrA-*S83Y | 48 (10.1%) | 0 (0.00%) | 48 (5.88%) |
| *bla_TEM-1_* *, catA1, dfrA7, sul1, sul2, strAB, tetB* / *gyrA-*S83F | 2 (0.42%) | 0 (0.00%) | 2 (0.25%) |
| *bla_TEM-1_* *, catA1, dfrA7, sul1, sul2, strAB, tetB* / *gyrA-*D87N | 1 (0.21%) | 0 (0.00%) | 1 (0.12%) |
| *bla_TEM-1_* *, catA1, dfrA7, sul1, sul2, strAB, tetB* / *gyrA-*S83Y, *parC-*S80R | 1 (0.21%) | 0 (0.00%) | 1 (0.12%) |
| *blaTEM-1 , catA1, dfrA7, sul1, sul2, strAB, qnrS, tetA /* gyrA-S83F | 1 (0.21%) | 0 (0.00%) | 1 (0.12%) |
| *blaTEM-1, catA1, dfrA7, sul1, strAB, tetB / gyrA-S83Y* | 1 (0.21%) | 0 (0.00%) | 1 (0.12%) |
| *bla_TEM-1_* *, catA1, dfrA7, sul1, sul2, strAB, tetB* | 1 (0.21%) | 0 (0.00%) | 1 (0.12%) |
| *blaTEM-1 , catA1, sul2, strAB / gyrA-S83Y* | 1 (0.21%) | 0 (0.00%) | 1 (0.12%) |
| *bla_TEM-1_* *, catA1, dfrA7, sul1/ gyrA-*S83F | 2 (0.42%) | 0 (0.00%) | 2 (0.25%) |
| *bla_TEM-1_* *, catA1, dfrA7, sul1 / gyrA-*S83Y | 6 (1.26%) | 0 (0.00%) | 6 (0.74%) |
| *catA1, dfrA7, sul, tetB /* gyrA-S83Y | 1 (0.21%) | 0 (0.00%) | 1 (0.12%) |
| *catA1,sul2, strA, strB /* gyrA-S83F | 1 (0.21%) | 0 (0.00%) | 1 (0.12%) |
| *catA1, dfrA7, sul1/ gyrA-*S83F | 47 (9.81%) | 0 (0.00%) | 47 (5.76%) |
| *catA1, dfrA7, sul1/ gyrA-*D87N | 2 (0.42%) | 0 (0.00%) | 2 (0.25%) |
| *catA1, dfrA7, sul1/ gyrA-*S83F, *gyrA-D87G, parC-*E84K | 1 (0.21%) | 0 (0.00%) | 1 (0.12%) |
| *catA1, dfrA7, sul1* | 1 (0.21%) | 0 (0.00%) | 1 (0.12%) |
| *bla_TEM-1_, sul2, qnrS, tetA /* *gyrA-*S83Y | 62 (12.94%) | 0 (0.00%) | 62 (7.59%) |
| *sul1, strAB, tetB / gyrA-*S83Y | 1 (0.21%) | 0 (0.00%) | 1 (0.12%) |
| *bla_TEM-1_, sul2, qnrS / gyrA-*S83Y | 5 (1.05%) | 0 (0.00%) | 5 (0.61%) |
| *bla_TEM-1,_ bla_CTX-_*_M_ | 0 (0.00%) | 1 (0.29%) | 1 (0.12%) |
| **Only QRDR mutation** | **51 (10.6%)** | **248 (73.16%)** | **299 (36.55%)** |
| *gyrA-*S83F | 19 (3.98%) | 190 (56.05%) | 209 (25.6%) |
| *gyrA-*S83Y | 27 (5.66%) | 13 (3.83%) | 40 (4.9%) |
| *gyrA-D87G* | 2 (0.42%) | 1 (0.29%) | 3 (0.37%) |
| *gyrA-D87N* | 0 (0.00%) | 37 (10.91%) | 37 (4.52%) |
| *gyrA-D87Y* | 2 (0.42%) | 6 (1.77%) | 8 (0.98%) |
| *gyrA-*S83F, *parC-*E84K | 0 (0.00%) | 1 (0.29%) | 1 (0.12%) |
| *gyrA-*S83F, *gyrA-*D87N, *parC-*S80I | 1 (0.21%) | 0 (0.00%) | 1 (0.12%) |
| **Susceptible to all antibiotics** | **6 (1.25%)** | **90 (26.55%)** | **96 (11.74%)** |
